# Urbanization and species-energy relationships

**DOI:** 10.1101/431189

**Authors:** Lucas M. Leveau

## Abstract

Species-energy relationships (SER) have been used scarcely to predict responses of biodiversity to changes in land use. In this work, two hypothesis of the SER were tested on bird communities along urbanization gradients: the increasing population size (IPS) and the niche position (NP) hypotheses. The IPS hypothesis states that sites with more primary productivity have more individuals, decreasing the probability of extinction and increasing species richness. The HPN predicts that sites with greater primary productivity and environmental heterogeneity allow the availability of a greater variety of resources and the presence of specialist species, thus increasing species richness. The hypothesis of habitat heterogeneity was evaluated by which more complex habitats provide more variety of resources and greater species richness. Birds were counted along three urban-rural gradients in the Pampean region of Argentina during the breeding season. The productive energy was measured from the NDVI and the environmental heterogeneity through the Shannon-Wiener diversity index, calculated with the percentage coverage of different substrates. Bird richness and habitat diversity were higher at intermediate levels of urbanization, while primary productivity had a negative relationship with urbanization. Although there was a positive relationship between bird richness and abundance, there was no positive relationship between bird abundance and primary productivity. Bird richness showed a positive relationship with habitat diversity and primary productivity; although path analysis revealed that the habitat diversity increased bird richness through an increase of bird abundance. The NP hypothesis was the most consistent to predict the bird richness along urban-rural gradients.

## 1. Introduction

Species-energy relationships have been increasingly invoked over the last three decades to explain the spatial and temporal variation of species richness (Wright 1983, Rosenzweig 1995, Evans et al. 2005). Depending on the spatial scale and taxa, species richness may increase monotonically with energy, or may initially increase but then decline at higher energy levels in a unimodal relationship (Mittelbach et al. 2001, Pärtel et al. 2007, Fraser et al. 2015). Despite a large number of studies published about the species-energy relationship, there is a need to understand the mechanism underlain such relationship. Evans et al. (2005) identified nine general mechanisms that may generate positive species-energy relationships at the macroscale. However, several mechanisms such as the increasing population size (IPS) and the niche position (NP) hypothesis may be tested at smaller scales.

Although the species-energy relationships have been extensively studied in natural environments (Tellería et al. 1992, Srivastava and Lawton 1998, Honkanen et al. 2010), there are few studies in heavily impacted environments. One of the most extreme human alterations on Earth is the urbanization of natural and rural areas. Through habitat loss and fragmentation, pollution and altered resource flow, urbanization promotes a biodiversity loss (Grimm et al. 2008, Shanahan et al. 2014). On the other hand, urbanization is thought to increase animal abundance through the increase of subsidized food by humans and a decrease of predation (Shochat et al. 2006, but see Saari et al. 2016). Therefore, lower species richness and a higher abundance in urban environments disagree with the IPS mechanism, which states that higher population sizes promote higher species richness by reducing extinction risk (Srivastava and Lawton 1998, Forsman and Mönkkönen 2003, Evans et al. 2005). However, Chiari et al. (2010) tested this mechanism along the urban gradient of Florence (Italy) finding a positive species-abundance relationship.

Habitat heterogeneity and energy are potential indicators of the niche position hypothesis (Evans et al. 2005, Hurlbert 2004). More energy availability and higher habitat heterogeneity are indicators of more resource types that are above the abundance threshold at which they can support a viable population (see Figure 2 in Evans et al. 2005). Therefore, there can be a higher number of niche position specialists in sites with high habitat complexity and high energy availability, elevating the species richness compared to sites with low habitat heterogeneity and energy availability (Hurlbert 2004, Honkanen et al. 2010, Elo et al. 2012). Moreover, as the number and quantity of resources increases, they are divided more equitably among a greater variety of foraging habitats; then, a lower dominance is expected as energy availability and habitat diversity increase (Hurlbert 2004). Due to this increase of new resource types changes along the gradient, a shift of species composition is expected because generalist species are replaced by specialist species (Harrower et al. 2017).

The habitat heterogeneity hypothesis (HH) is another fundamental mechanism explaining the spatial variation of bird richness (MacArthur 1964, Hurlbert and Haskell 2003, Tews et al. 2004). A more variety of vegetation types is related to a greater quantity of different ways to obtain resources, resulting in a higher coexistence of species (Rosenzweig 1995). But the importance of habitat heterogeneity determining the species richness in urban environments has been scarcely studied. Intermediate levels of urbanization characterized by detached houses with yards are generally composed by different layers of vegetation favouring higher habitat diversity than rural or natural areas, especially for arid shrubland and grassland biomes (McKinney 2002, Miller et al. 2001, Leveau 2013a). This, in turn, may favour higher animal species richness in suburban areas, forming a unimodal species richness-urbanization relationship (Blair 1996, Leveau and Leveau 2005, Tratalos et al. 2007).

The aim of this study is to evaluate which mechanism is more likely explaining the bird species richness variation along three urban-rural gradients of the Pampean region, Argentina. Although several studies have proposed theoretical models of bird richness response to urbanization (Lepczyk et al. 2008, Filloy et al. 2018), species-energy relationships have been scarcely studied and, to my knowledge, several hypotheses have never been explored simultaneously in urban environments yet. Unlike natural systems, in which primary productivity and habitat diversity are usually correlated (Hurlbert 2004), urban-rural gradients in central Argentina allows to separate the effects of these variables because agricultural areas can have a high primary productivity but a low habitat heterogeneity due to crops dominance (Leveau 2013a, Leveau et al. 2018). Due to bird richness has been shown an unimodal relationship with urbanization in the study area (Leveau and Leveau 2004, 2005), increasing at intermediate levels of urbanization that have both high primary productivity and habitat diversity (Leveau 2013a, Leveau et al. 2018), I expect that the NP hypothesis will explain more clearly the spatial variation of bird richness in central Argentina. On the other hand, Path analyses is a powerful statistical approach in testing and developing the structural hypothesis with both indirect and direct causal effects (Fan et al. 2016), which has been used in studies of species-energy relations (Jonsson et al. 2011, Elo et al. 2012, McCain et al. 2018). Therefore, a Path analysis was performed to analyze the direct and indirect effects of primary productivity, habitat diversity and bird abundance on bird richness.

## 2. Methods

### 2.1. Study area

The study was made in three cities of the Pampean region, in Buenos Aires province (Argentina): Mar del Plata (615 350 inhabitants), Balcarce (38 823 inhabitants) and Miramar (29 629 inhabitants). The city sizes in this study controls for the city-size effect found in other studies, focusing on cities of more than 20 000 inhabitants (Jokimäki and Kaisanlahti-Jokimäki 2003, Garaffa et al. 2009). The maximum distance between the cities in Argentina was 59 km (between Balcarce and Miramar), thus the effects of latitude or climate are minimum. These cities are surrounded by a landscape composed of cultivated land, pastureland, grasslands and exotic tree plantations. The climate is temperate and the maximum mean temperature is 20.3° C during January and the mean annual precipitation is 923.6 mm (data from the Meteorological National Service).

### 2.2. Bird census

Bird surveys were made in three habitat types: 1) urban centre; 2) suburban areas composed of detached houses with yards; and 3) rural areas, composed by crops and pastures (Figure 1). In each habitat I placed five transects of 100 × 50 m separated each other by at least 200 m. During each breeding season, corresponding to spring-summer of the years 2011–2012 and 2012–2013, transects were visited twice totalling four visits to each transect. Surveys were conducted in the first 4 h after dawn on days without rain or strong winds, and all birds seen or heard were counted, except those flying over the top of buildings or trees or below that height but without feeding activity.

**Figure 1.**
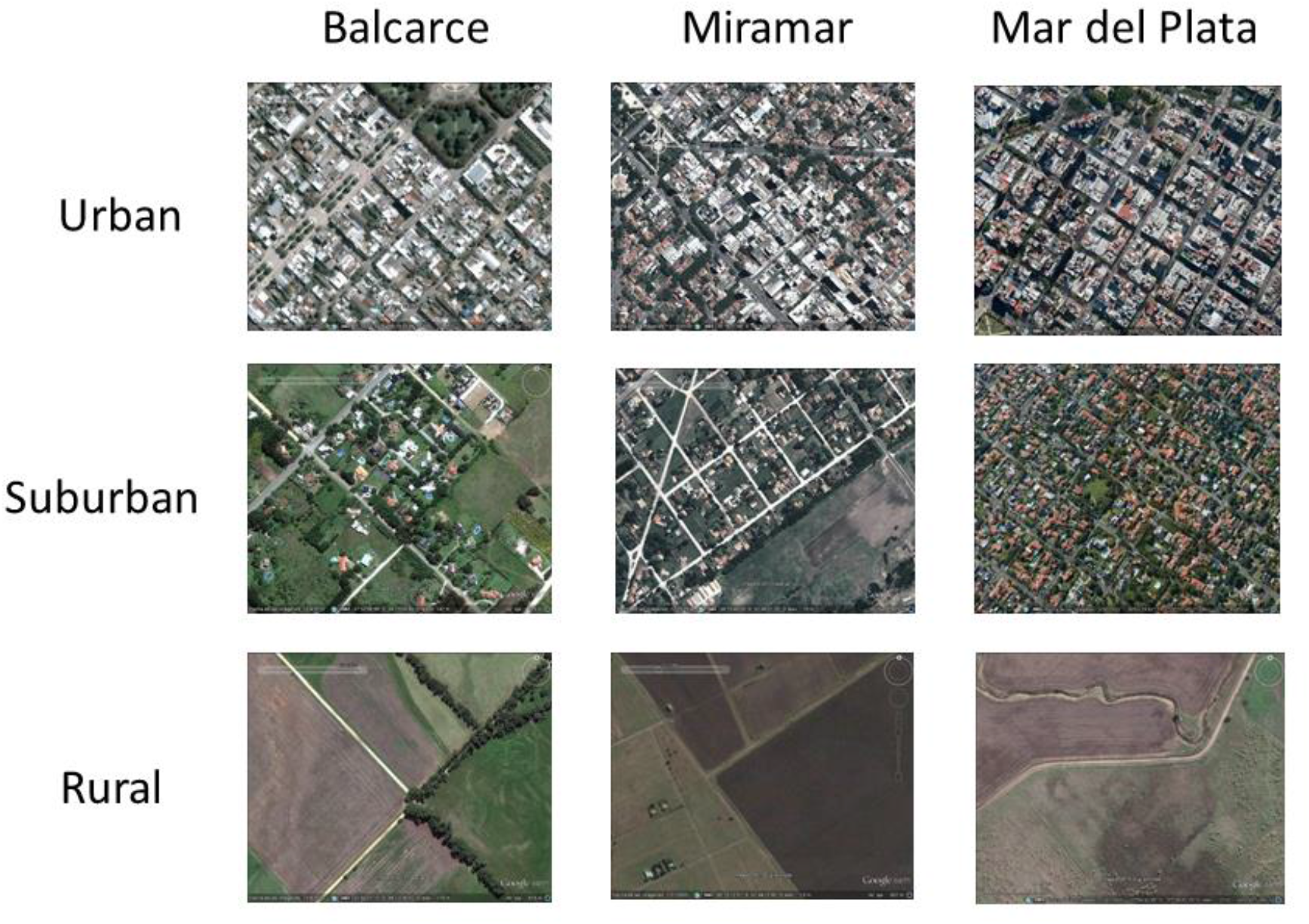
Study design showing Google earth images of surveyed habitat types (rows) along urban-rural gradients of three cities (columns) in central Argentina.

### 2.3 Urbanization, habitat diversity and primary productivity

In each transect, two circles of 25 m radius were located, one in the centre of the first 50 m along transects and the other in the centre of the remaining 50 m. In each circle, the following variables were estimated visually: 1) percentage coverage of trees, 2) shrubs, 3) lawn (managed herbaceous vegetation), 4) buildings, 5) non-managed herbaceous vegetation, 6) cultivated land, and 7) paved roads. We summed the percentage cover of buildings and paved roads to get a measure of impervious surface cover in each circle and averaged values of both circles in each transect. On the other hand, habitat diversity in each transect was estimated using the Shannon index, which incorporated the percent cover of trees, shrubs, lawn, herbaceous vegetation, cultivated land and buildings. When the percentage cover of habitat components exceeded 100 %, values were corrected for up to 100 %.

For primary productivity I used the Normalized Difference Vegetation Index (NDVI), which is a measure of greenness that correlates positively with the net primary productivity (Rouse et al. 1974; Pettorelli et al. 2005) and is assumed to correlate positively with production of the food available to birds (Hurlbert and Haskell 2003). The product MOD13Q1 from the MODerate resolution Imaging Spectroradiometer (MODIS sensor; Terra satellite) was used (Justice et al. 2002), available in https://lpdaac.usgs.gov/products/modis_products_table, which provides NDVI values in periods of 16 days with a spatial resolution of 250 m (Huete et al. 2002). For each period, an algorithm applies a filter to the data based on quality, cloud and viewing geometry. Then, the best observation of each pixel is selected, which represented the NDVI value of that period. The MODIS products are distributed according to tiles, which have 1200 × 1200 km in a sinusoidal projection. In this study, the h13v12 tile images were downloaded. Images encompassing the first and second visit of each breeding season were downloaded, and NDVI values were averaged for each transect. Images were referenced using the MODIS reprojection Tool software, (https://lpdaac.usgs.gov/tools/modis_reprojection_tool), using the UTM projection. Then, the images were reprojected again to the Argentina Gauss-Krüger projection.

Although the transect sizes (0.5 ha) were much smaller than the pixel sizes (6.25 ha), it was assumed that the NDVI values obtained for each transect represent the resources available for birds. On the other hand, birds may use resources in areas larger than the transect size. Hostetler and Holling (2000) found that birds in urban environments may have sufficient resources at areas of 0.2 km^2^ to 1.5 km^2^. Bino et al. (2008) found that bird richness had the best fit to NDVI when it was calculated at a pixel size of 14.8 ha, higher than their sampling unit size (point counts of 50 m radius), suggesting that the sensor MODIS may be efficient in depicting resources available for birds.

### 2.4 Statistical analysis

First, I wanted to explore the relationship between impervious cover (paved roads and buildings) and the primary productivity and habitat diversity. Due to the possible presence of non-linear relationships among variables, Generalized Additive Models (GAMs) with a Gaussian distribution of errors were conducted using the mgcv package of R (Wood 2011, R Development Core Team 2011).

Bird richness in each transect was calculated using a capture-recapture approach which assumes heterogeneity in detection probability among species and sites (Burnham and Overton 1979; Boulinier et al. 1998). This procedure uses a jackknife estimator associated with model M(h) (Burnham and Overton 1978, 1979; Otis et al. 1978), which estimates bird species richness and species detection probability. Therefore, we took into account possible differences in detectability among species within a transect and possible differences in detectability among transects. These estimates were computed using COMDYN software (Hines et al. 1999), with presense-absence data of species in the four visits to each transect. On the other hand, bird abundance was estimated as the mean number of birds recorded during the four visits to each transect. Bird dominance was calculated as the proportion of bird abundance of the two most abundant species over the total of bird species abundances during each of the four visits to the sampling units; an average of bird dominance over the four visits was calculated.

I tested three mechanisms that would explain the bird richness variation along the three urban-rural gradients. First, I evaluated the IPS hypothesis, where I expected a direct relationship between bird richness and abundance and a direct relationship between bird abundance and primary productivity. Second, the NP hypothesis was evaluated, where an increase in bird richness and a decrease in bird dominance was expected in sites with high primary productivity and habitat diversity. Moreover, a significant change in species composition was expected along the gradient of primary productivity and habitat diversity. Finally, I tested the HH hypothesis, where a direct relationship between bird richness and habitat diversity was expected. Because NP and HH models were nested, a Likelihood ratio test (LRT) was performed among them to test the significance of primary productivity explaining the variation of bird richness along the gradients.

Due to the nested design of the study, in which transects were nested in cities, Generalized Linear Mixed Models (GLMMs) were conducted to test the different mechanism explaining bird richness along the urban gradients. Bird richness data were log transformed to improve normality and homoscedasticity of residuals. The city was deemed as a random factor, and the package nmle in R was used. The significance of the nested models was assessed by comparing them with Generalized least squares (GLSs) models through an LRT. In all cases, the random effect was not significant (P > 0.05); therefore, I conducted GLSs. The correlation between primary productivity and habitat diversity was low (Pearson coefficient, r = 0.31).

Bird species composition was analyzed by a Non-metric multidimensional scaling (NMDS). A matrix of bird species as columns and bird abundance in each sampling unit as rows was used to calculate a Bray-Curtis dissimilarity matrix. Then, an NDMS was performed and the values of primary productivity and habitat diversity were correlated to the resultant ordination by the function envfit of the package vegan in R (Oksanen et al. 2007). The arrow shows the direction of the environmental gradient, and the length of the arrow is proportional to the correlation between the environmental variable and the ordering of the NMDS (Oksanen 2015). Only significant vectors (P < 0.05) were plotted.

I explored direct and indirect effects of bird abundance, primary productivity and habitat diversity on bird richness through Path analysis (Grace and Pugesek 1998). The following relationships were tested: 1) bird richness was directly related to bird abundance, primary productivity and habitat diversity; 2) bird abundance was related to primary productivity and habitat diversity; and 3) primary productivity was related to habitat diversity. Models were adjusted by a backward selection with the anova function (see Appendix S1 in Grace et al. 2014), using the lavaan package in R (Rosseel 2012). The final model was selected through Chi-square tests (χ2), which try out the hypothesis that there is a discrepancy between the model-implied covariance matrix and the original covariance matrix (Fan et al. 2016). Then, the non-significant discrepancy is preferred (p > 0.05). The root mean square error of approximation (RMSEA) and standardized root mean square residual (SRMR) are “badness of fit” indices where 0 indicates the perfect fit and higher values indicate the lack of fit (Fan et al. 2016). The lavaan package performs a test which hypothesis is that RMSEA is less than or equal to 0.05, a cutoff sometimes used for “close” fit. A p-value greater than 0.05 does not reject the hypothesis and, therefore, the model has a good fit. The SRMR index should be less than 0.09 to be acceptable (Fan et al. 2016). Finally, the Comparative fit index (CFI) ranges between 0.0 and 1.0, and represents the amount of variance that has been accounted for in a covariance matrix; a value close to 0.95 or higher is acceptable (Fan et al. 2016). Bird abundance data was divided by 10 to have similar variance values to primary productivity and habitat diversity.

## 3. Results

A total of 54 species and 2200 individuals were counted (Table 1). Most abundant species were the House sparrow (*Passer domesticus*, 27 % of individuals), the Eared Dove (*Zenaida auriculata*, 14 %) and the Rock Dove (*Columba livia*, 12 %). The three habitat types have different bird composition. Urban centres were dominated by the Rock Dove, the Eared Dove and the House Sparrow, whereas suburban areas have more species and were dominated by the House Sparrow, the Eared Dove and the Picazuro Pigeon (*Patagioenas picazuro*). The rural areas were dominated by the Rufous-collared Sparrow (*Zonotrichia capensis*), the Grassland Yellow-finch (*Sicalis luteola*) and the House Sparrow.

**Table 1.**
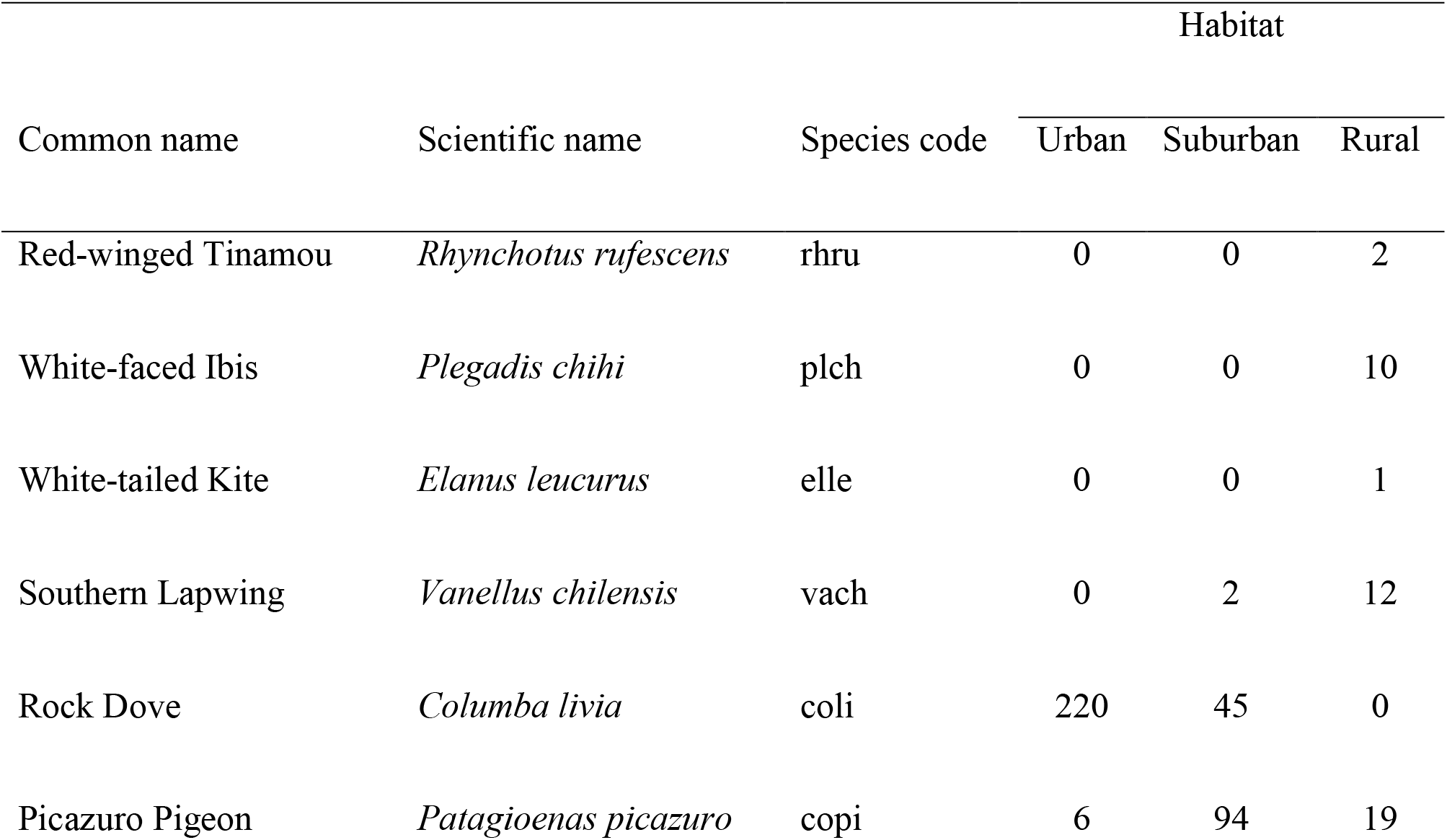
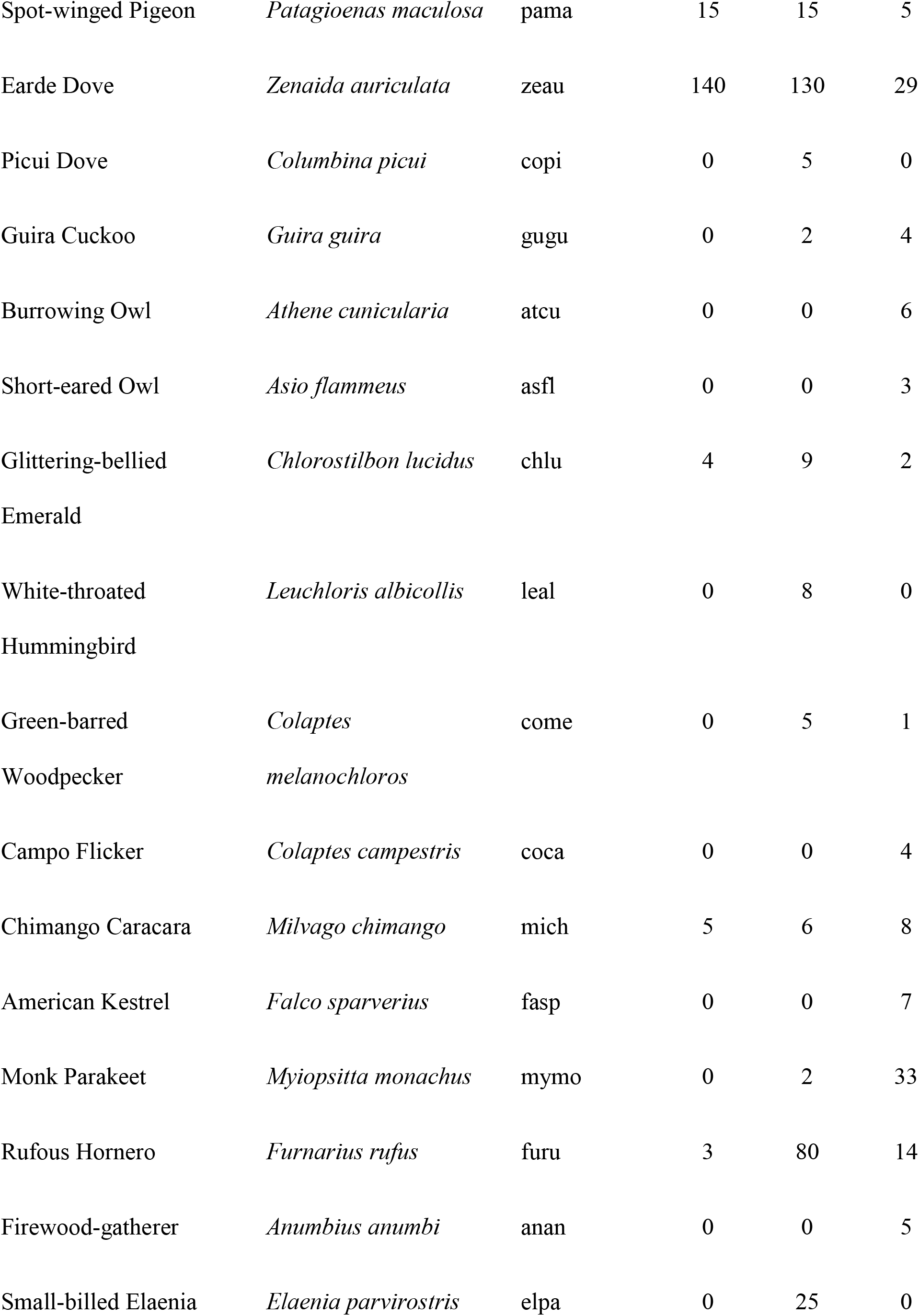
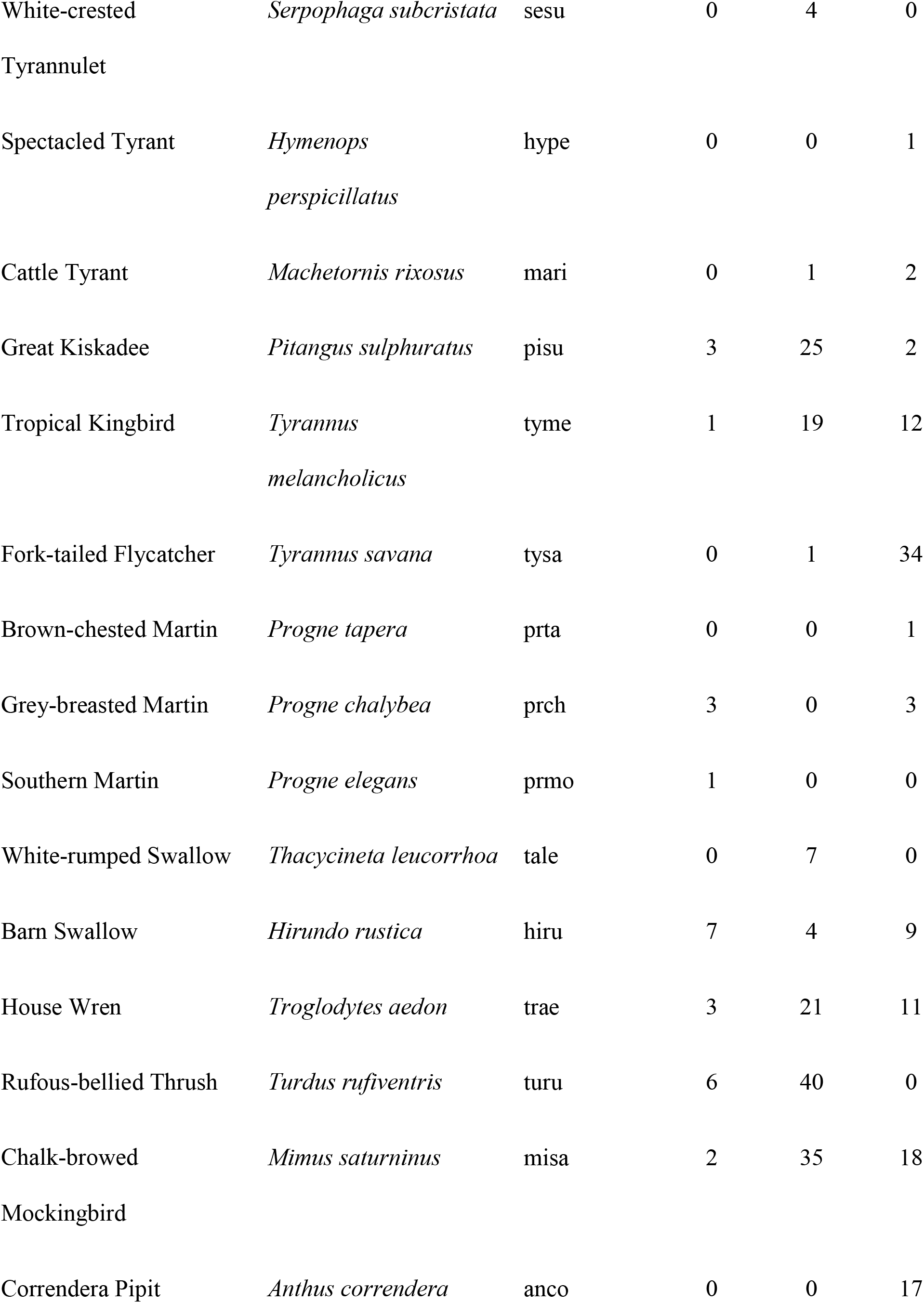
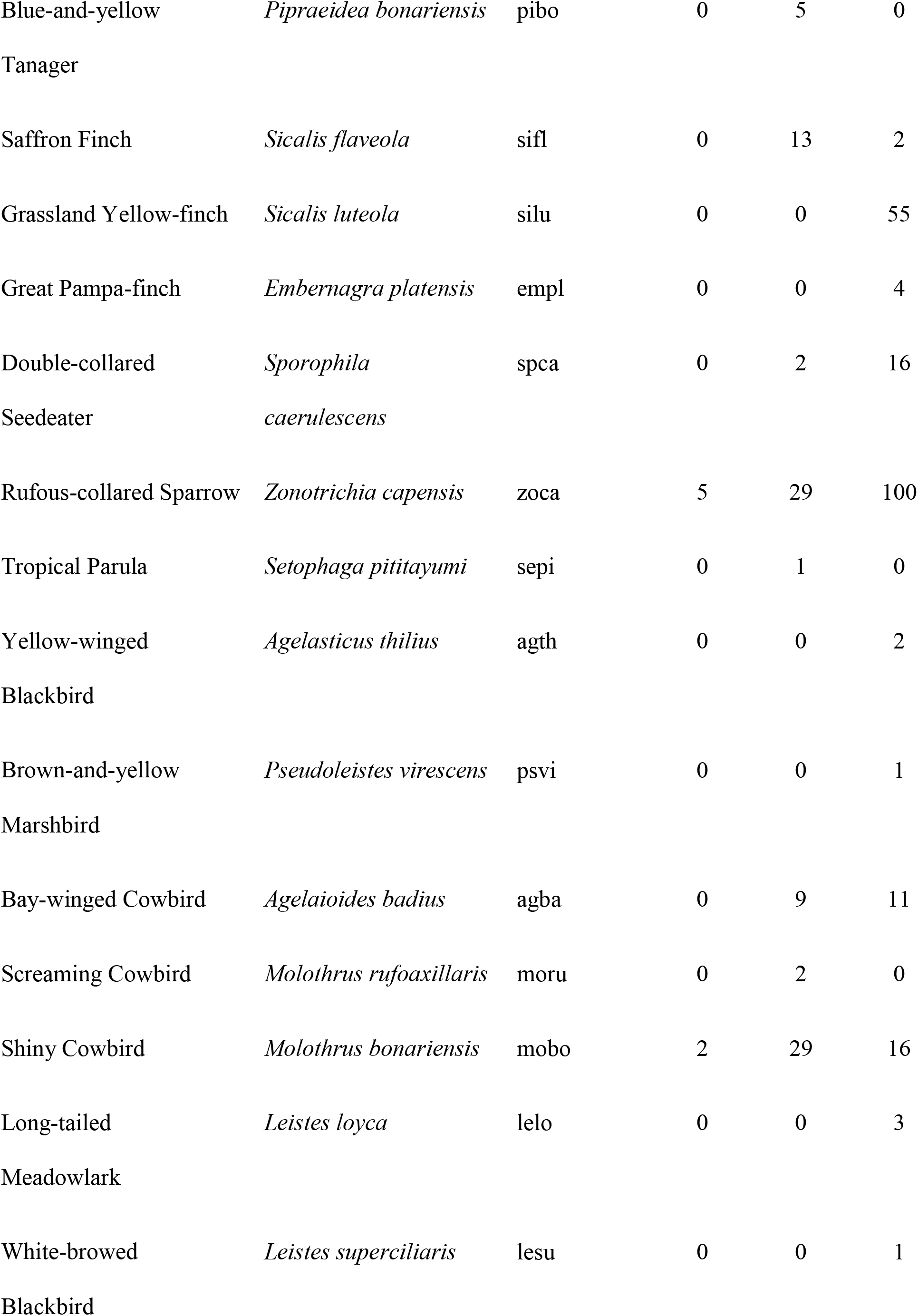
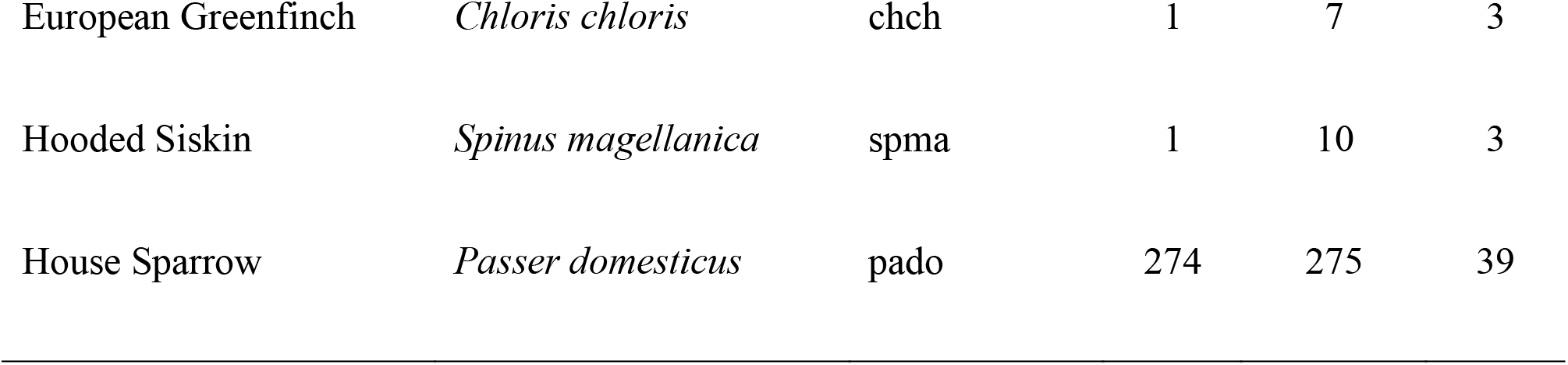
List of species observed in urban, suburban, and rural habitats of three cities in central Argentina.

Primary productivity declined with impervious surface cover (r^2^ = 0.88, F = 50.53, estimated degrees of freedom [edf]= 5.48, P < 0.001, n = 45; Figure 2), whereas habitat diversity was highest at intermediate levels of impervious surface (r^2^ = 0.53, F = 14.40, edf = 2.73, P < 0.001, n = 45). Therefore, energy availability was higher in rural areas whereas habitat complexity increased in suburban areas. Bird richness increased at intermediate levels of urbanization (r^2^ = 0.44, F = 12.55, edf = 2.32, P < 0.001, n = 45).

**Figure 2.**
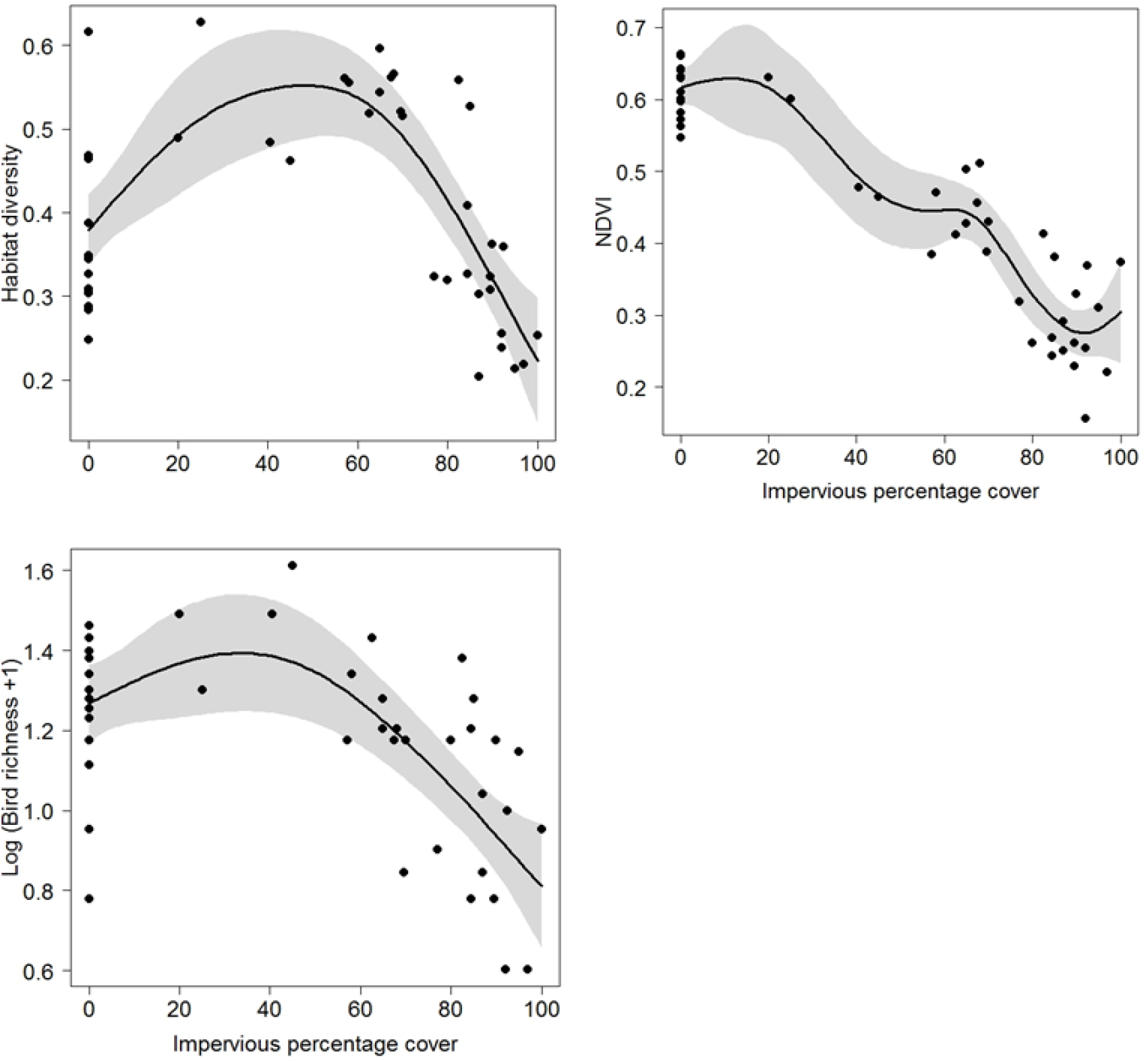
Relationships between habitat diversity (H′index), primary productivity (NDVI), bird richness (log) and impervious percentage cover in three cities of central Argentina. Models were constructed based on Generalized additive models.

Although bird species richness was related positively with bird abundance (Table 2), the latter was not related significantly with primary productivity (Figure 3). Therefore, the IPS hypothesis is not supported by results. On the other hand, although bird species richness had a positive relationship with habitat diversity supporting the HH mechanism (Table 2), the inclusion of primary productivity to the model had a better fit of data (LRT, F = 18.22, P < 0.001) (Figure 4). Moreover, dominance declined with higher primary productivity and habitat diversity (Figure 4), whereas species composition varied according to changes in habitat diversity and primary productivity along the gradients (Figure 5). Then, the NP mechanism would be the most likely explaining the variation of bird species richness along the urban gradients studied.

**Table 2.**
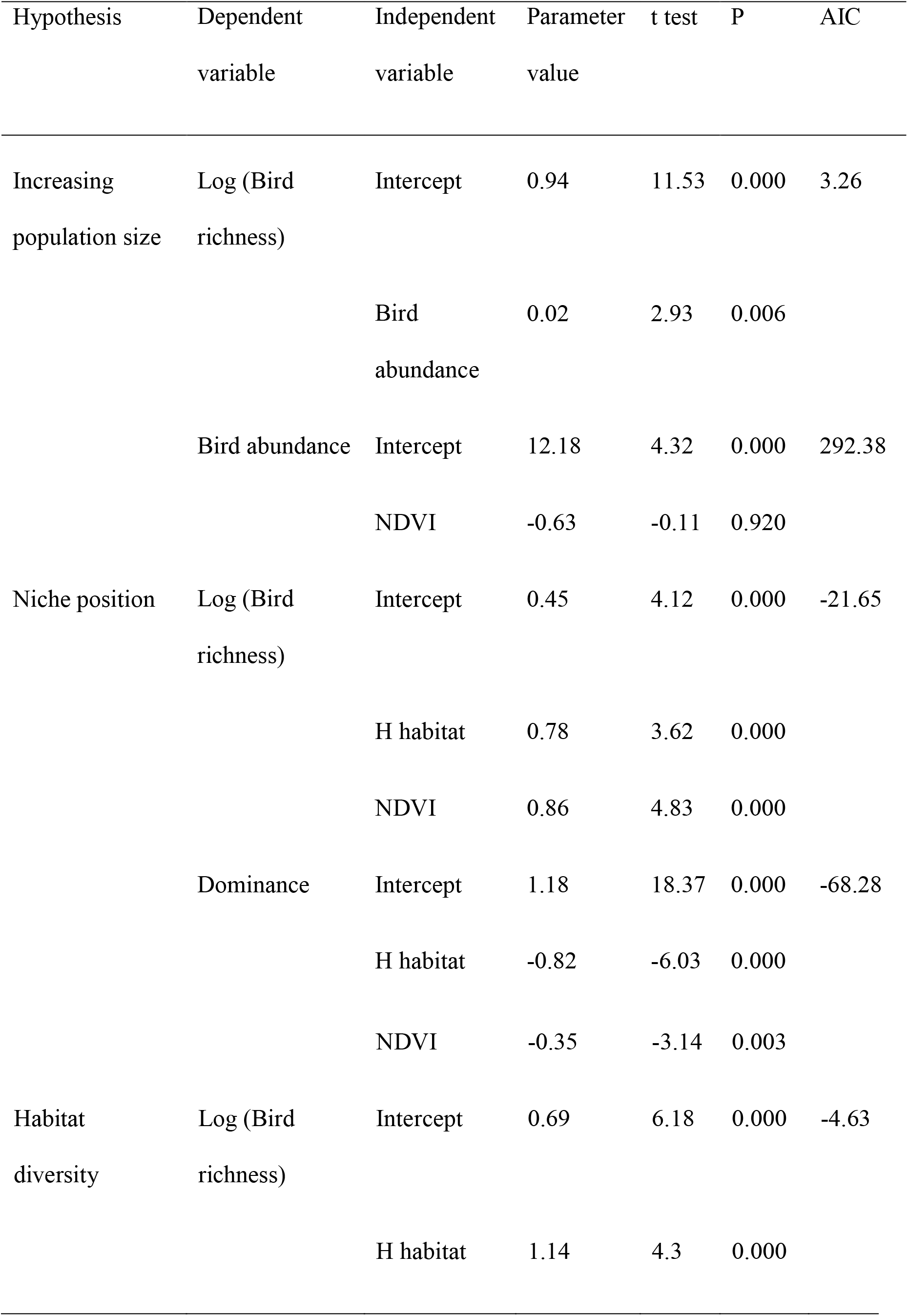
Main hypothesis and statistical models explaining the spatial variation of bird richness, abundance and dominance along urban-rural gradients of three cities of central Argentina.

**Figure 3.**
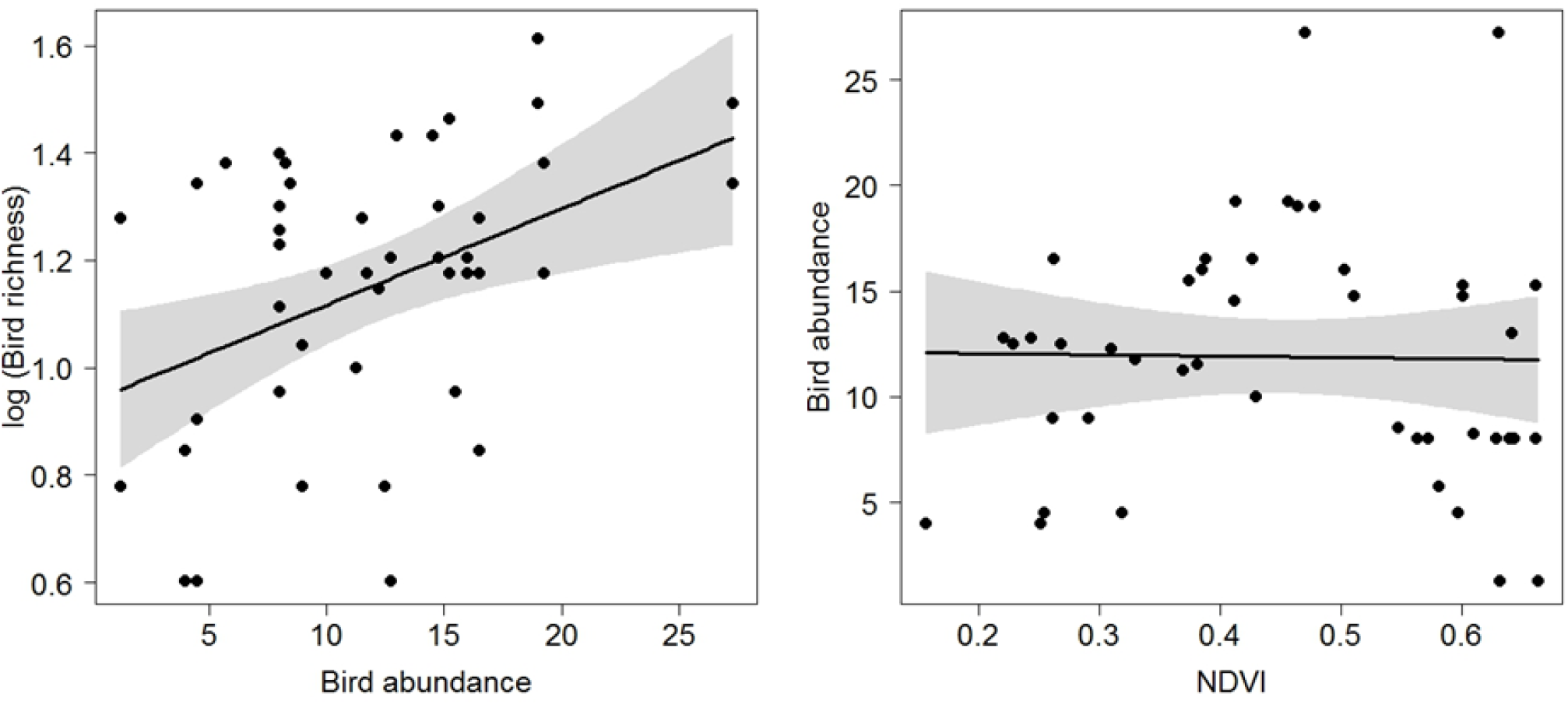
Relationships between bird richness and bird abundance and between bird abundance and primary productivity (NDVI) along urban-rural gradients of three cities in central Argentina.

**Figure 4.**
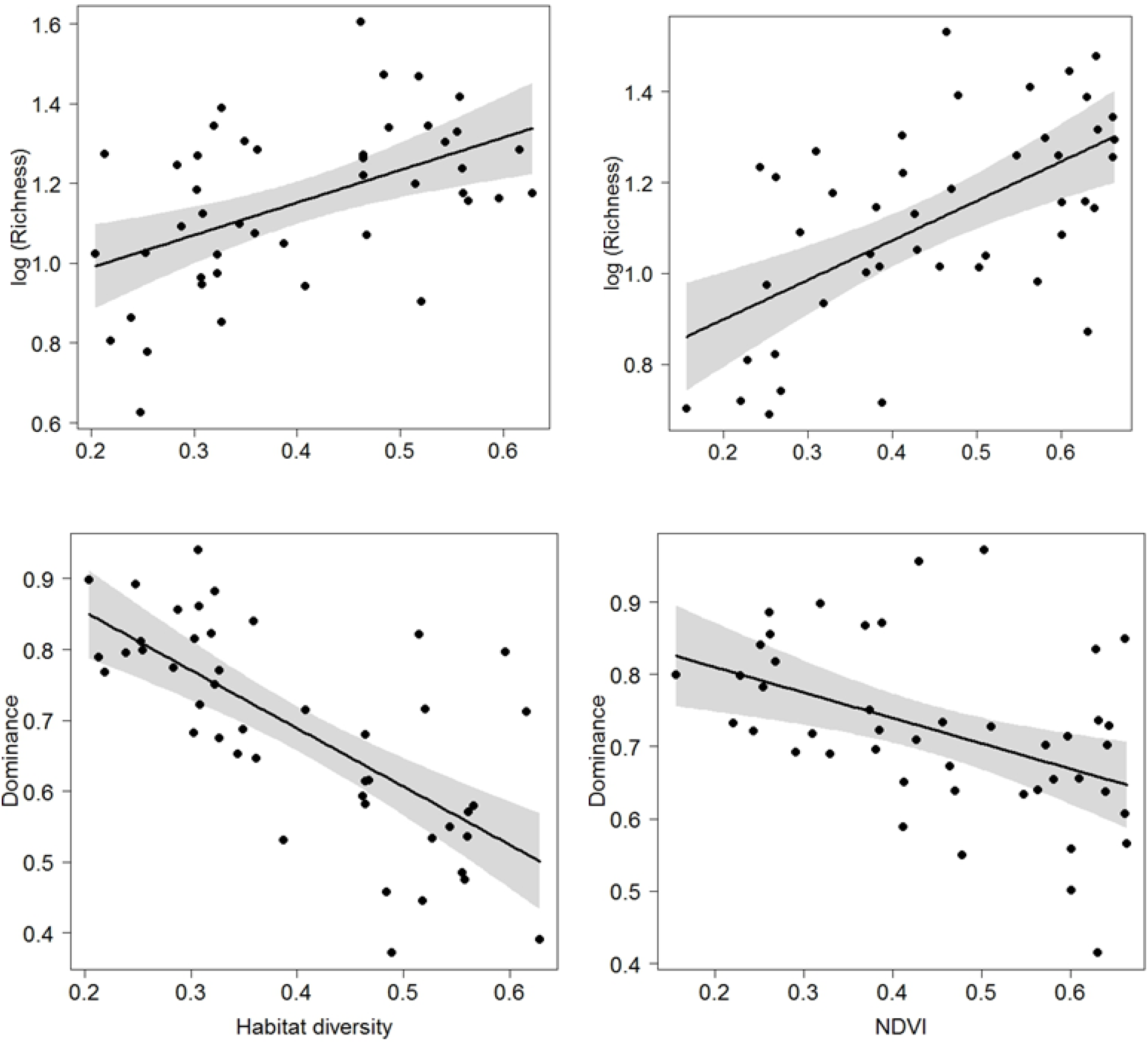
Relationships between bird richness, habitat diversity and primary productivity (NDVI), and relationships between bird dominance and habitat diversity and primary productivity along urban-rural gradients of three cities in central Argentina.

**Figure 5.**
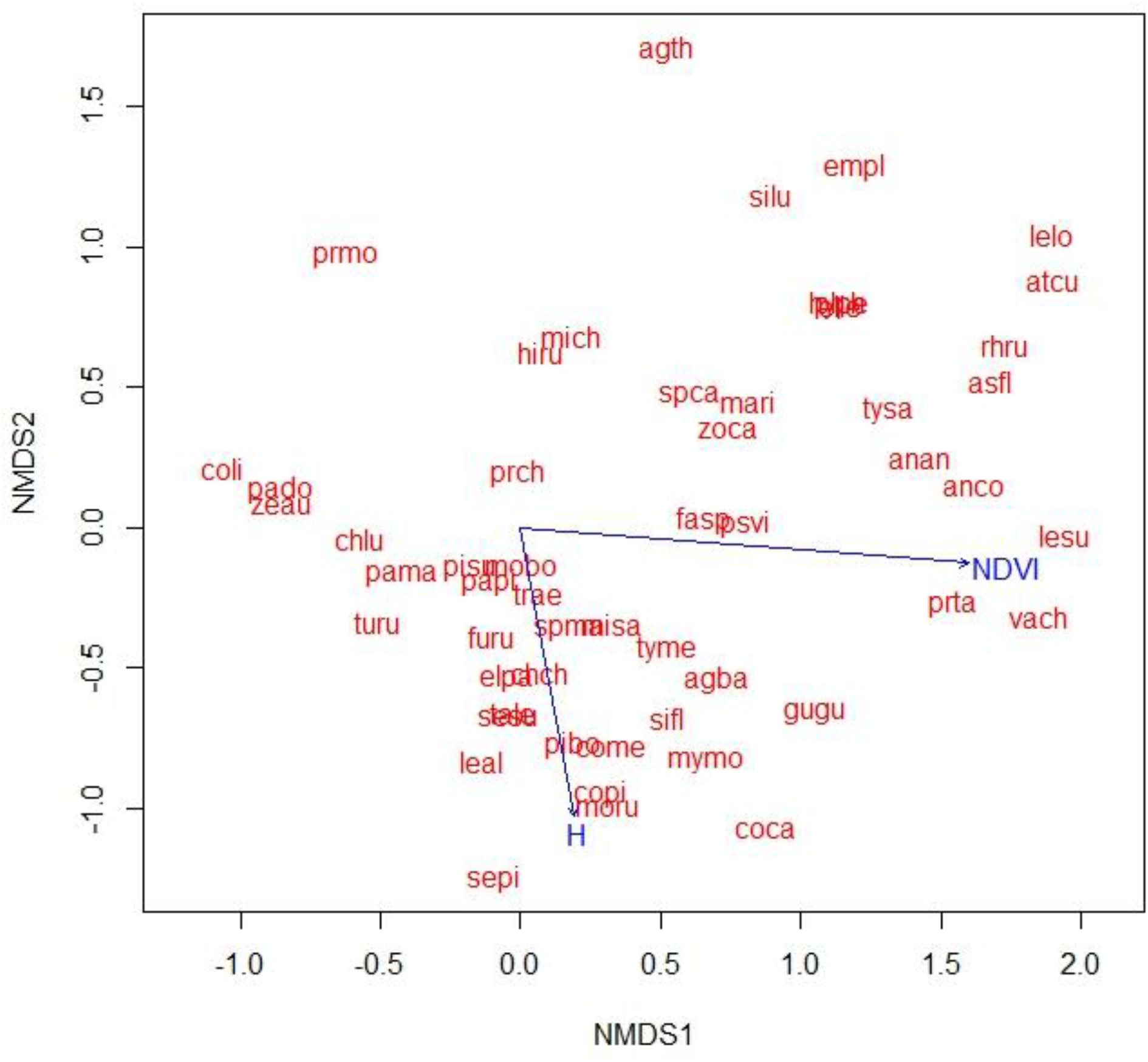
Non-metric multidimensional scaling showing the relationship between species composition along two axes and primary productivity (NDVI) and habitat diversity (H) along three urban-rural gradients in central Argentina. Stress = 0.18. For species names see codes in Table 1.

Path analysis had a good fit (χ^2^ test, p = 0.074; RMSEA = 0.19, p = 0.096; SRMR = 0.09; and CFI = 0.95). The model selected revealed direct relationships of primary productivity and bird abundance with species richness (Figure 6). Habitat diversity had a direct relationship with bird abundance and primary productivity, having an indirect effect on bird richness.

**Figure 6.**
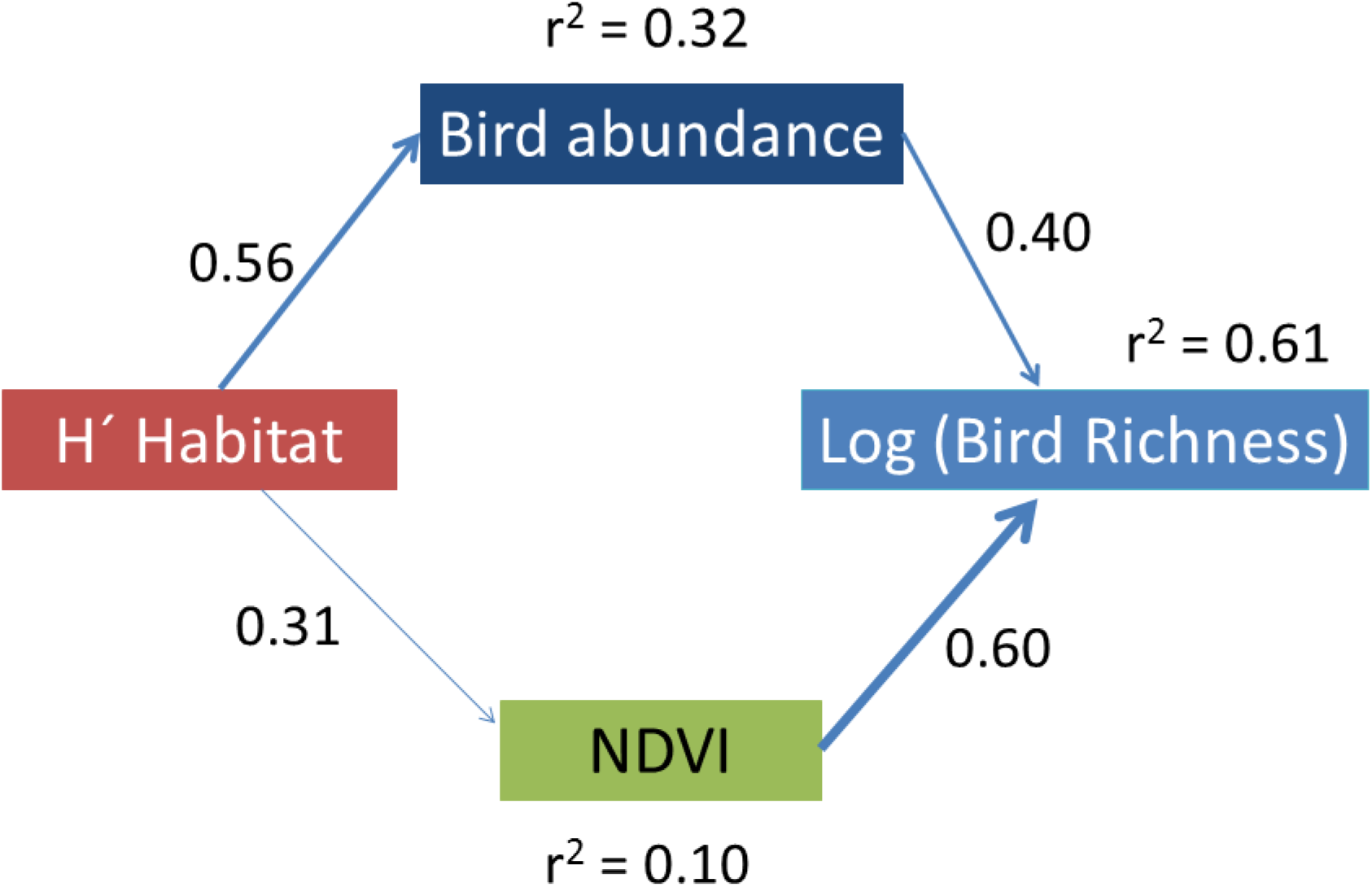
Path analysis showing the direct and indirect relationships between habitat diversity, primary productivity, bird abundance and bird richness. Standardized coefficients are showed (P < 0.05) with r-squared coefficients.

## 4. Discussion

Along the three urban-rural gradients analyzed in the Pampean region, urbanization affected negatively the primary productivity but increased the habitat heterogeneity at intermediate values of impervious surfaces. On the other hand, bird species richness was higher at sites with the highest primary productivity and habitat diversity.

Therefore, this result supports the NP hypothesis, which states that a more quantity and variety or resources allow the permanence of a higher number of species (Abrams 1995, Hurlbert 2004, Honkanen et al. 2010, Elo et al. 2012).

The suburban areas are the typical part of the urban-rural gradient where primary productivity and habitat diversity meets. Humans create yards with several vegetation strata: lawn, shrubs and trees. Usually, it is used a variety of shrub and tree species that provide a diversity of floral and fruit resources. The primary productivity is enhanced by the use of fertilizers and irrigation (Lepczyk et al. 2004). Buildings and trees may provide nesting places, allowing a higher species richness than highly urbanized areas and rural areas. As a result, suburban areas not only were inhabited by species that have a generalist diet and nest in buildings, such as the Rock Dove and the House Sparrow, but also by species that nest in trees and have an insectivorous diet, such as the Rufous Hornero (*Furnarius rufus*) and the Chalk-browed Mockingbird (*Mimus saturninus*), or a nectarivorous diet, as in the case of the White-throated Hummingbird (*Leucochloris albicollis*) and the Glittering-bellied Emerald (*Chlorostilbon lucidus*).

Highly urbanized areas had both lower habitat diversity and lower primary productivity, as a result of a dominance of impervious cover. However, they are inhabited by a few species (the Rock Dove, the Eared Dove and the House Sparrow) that have high abundances (see Table 2). These species may exploit food resources not resultant of primary productivity but instead provided by humans, such as food discards.

Rural areas had the highest primary productivity but lower habitat diversity than suburban areas, as a result of the dominance of crops. Therefore, sites were occupied mainly by granivorous species that nest on the ground, such as the Rufous-collared Sparrow and the Grassland Yellow-finch. However, an increase of habitat heterogeneity by the planting of trees or the presence of houses may increase species richness by allowing the presence of species that nest in trees, such as the the Moonk Parakeet (*Myiopsitta monachus*) and the Fork-tailed Flycatcher (*Tyrannus savanna*), or species that nest in holes, such as the House Sparrow or the Grey-breasted Martin (*Progne chalybea*).

Although the MIH has been postulated as a mechanism explaining bird richness variation along urbanization gradients (Chiari et al. 2009), our results showed a lack of relationship between primary productivity and bird abundance. Storch et al. (2018), based on a review, have also found a lack of support to the IPS for the most of their studies analyzed. Urban areas are characterized by a number of conditions that decouple the positive relation between bird abundance and bird richness. For example, the presence of constant food waste provided by humans, nest site availability and lower predation pressure enable to a reduced number of species to thrive in highly urbanized areas (Shochat et al. 2006, Shanahan et al. 2014).

Path analysis revealed that the effect of habitat diversity on bird richness was not direct, but rather mainly through increasing bird abundance. The positive effect of habitat diversity on bird abundance has been shown to be mainly direct on bird richness (Elo et al. 2012). Our results highlight the importance to analyze what is the role of habitat diversity in the niche position hypothesis; whether its effect is direct on bird richness or indirect by increasing bird abundance. Habitat diversity may favour greater bird abundance by providing more refugees (Kohn and Leviten 1976). Alternatively, the presence of different overlapped vegetation layers may provide more volume that can hold more bird individuals per habitat area (Dean and Connell 1987). A greater number of individuals may increase species richness by a sampling phenomenon (O’Connor 1991).

### 4.1 Conservation implications

The three processes tested have different management decisions to increase species richness along urbanization gradients (Harrower et al. 2017). If the HH hypothesis was supported, only efforts to create more diverse habitats would be adequate. If the IPS hypothesis was verified, a focus on more vegetation plantation would be good to allow higher bird richness. However, our results supported the NP hypothesis and, therefore, habitats with both high vegetation cover and a high habitat complexity would be necessary to allow a higher species richness. However, in our study area, sites with higher species richness can be inhabited by species with low conservation status that are widely distributed in the region, such as the Rufous-bellied Thrush (*Turdus rufiventris*), the Great Kiskadee (*Pitangus sulphuratus*) or the Rufous Hornero (*Furnarius rufus*). On the other hand, the NP hypothesis is consistent with a gradual change of species composition along the disturbance gradient (Harrower et al. 2017). Indeed, our data showed that bird communities along the urban-rural gradient were dominated by different species (Table 2). Therefore, conservation efforts also must focus on the specific habitat requirements of individual species. Although rural areas may have lower species richness than suburban areas, they are inhabited by species typical of grasslands that nest on the ground, such as the Short-eared Owl (*Asio flammeus*), the Grassland Yellow-finch (*Sicalis luteola*) and the Great Pampa-finch (*Embernagra platensis*).

It is evident that conservation implications are dependent on the scale of the study. If the focus is to promote bird richness within the city, more emphasis should be done to create areas with several types of vegetation. On the other hand, if the focus is on the consequences of urban expansion over rural areas, more importance should be put on the conservation of species typical of grasslands.

## 5. Conclusions

Although several studies have shown the importance of primary productivity (Haedo et al. 2017, Leveau et al. 2018, Filloy et al. 2018) and habitat diversity (Leveau 2013b) determining the spatial variation of species richness in urban areas, this is the first case study where both variables are considered in the context of species-energy relationships. The niche position hypothesis was supported and bird richness increased with more primary productivity and habitat diversity. However, the effect of habitat diversity was indirect through an increase of bird abundance. For a conservation point of view, this implies that focusing on quantity and variety of vegetation is the best option to enhance bird richness, but particular conservation efforts must be done to conserve grassland species which have specific habitat requirements and are negatively affected by the urban expansion.

